# Learning sequence-based regulatory dynamics in single-cell genomics

**DOI:** 10.1101/2024.08.07.605876

**Authors:** Ignacio L. Ibarra, Johanna Schneeberger, Ege Erdogan, Lennart Redl, Laura Martens, Dominik Klein, Hananeh Aliee, Fabian J. Theis

**Affiliations:** Institute of Computational Biology, Helmholtz Center Munich, Germany; Department of Mathematics, Technical University of Munich, Germany; TUM School of Life Sciences Weihenstephan, Technical University of Munich, Germany

## Abstract

Epigenomics assays, such as chromatin accessibility, can identify DNA-sequence-specific regulatory factors. Models that predict read counts from sequence features can explain cell-based readouts using specific DNA patterns (genomic motifs) but do not encode the changes in genomic regulation over time, which is crucial for understanding biological events during cell transitions.

To bridge this gap, we present *muBind*, a deep learning model that accurately predicts genomic counts of single-cell datasets based on DNA sequence features, their cell-based activities, and cell relationships (graphs) in a single architecture, enhancing the interpretability of cell transitions due to the possibility of inspecting motif activities weighted by nearest neighbors.

*MuBind* shows competitive performance in bulk and single-cell genomics. When complemented with graphs learned from RNA-based dynamical models used as injected priors in our model, *muBind* enhances through motif-graph interactions the identification of transcriptional regulators explaining cell transition events, including Sox9 in pancreatic endocrinogenesis scATAC-seq, and Gli3/Prdm16 in mouse neurogenesis and human organoids scRNA-seq, both supported by independent evidence, including associations between chromatin and motif activities over pseudotime, TF-gene expression patterns, and biological knowledge of these regulators.

*muBind* advances our understanding of cell transitions by revealing regulatory motifs and their interactions, providing valuable insights for genomic research and gene regulatory network dynamics. It is available at https://github.com/theislab/mubind.

## Introduction

Understanding the genomic control of cellular processes is key to defining cell identity and differentiation. Phenotypic variation is encoded in DNA sequences, regulated by transcription factors (TFs)^1^. Genomic measurements, such as chromatin accessibility (ATAC-seq), histone modifications, and DNA methylation^2–4^ profile cells and elucidate these rules.

Computationally, identifying TF rules involves recognizing DNA sequence patterns predictive of genomic observations like read counts^5^. Position-weight matrices (PWMs)^6^ summarize nucleotide-level contributions and identify short DNA patterns (motifs) associated with direct TF-DNA interactions^7^, as well as artifacts^8,9^. Deep learning (DL) models, such as convolutional neural networks and transformers^10,11^, improve performance and scalability. An alternative modeling strategy uses learnable convolutional filters and iterative, greedy optimization for faster convergence, allowing direct interpretation of main signal contributors^12^, yet potentially affecting generalization performance in scalable applications due to constraining the learning task to a limited set of motifs^13^. Current DNA sequence-based models treat samples as independent entities, or as species/cell-type groups, and miss cell connectivities essential to understanding cellular processes in pseudotemporal representations^14^, able to be generated with single-cell computational methods^15^. Graph-based representations of cell dynamics, like *k*-nearest neighbor (kNN) graphs and RNA velocity, refine sequence-based models by leveraging cell relationships. These can be complemented with cell dynamics information, like graph-based representations of RNA unspliced and spliced read counts, and transition matrices^15^. This opens up the possibility to refine sequence-based activities of a sequence model through operations that consider cell relationships. *ProBound*^*12*^ propagates sequence-inferred activities across bulk samples with experimentally designed relationships, yet single-cell genomics applications require flexible graphs representations. Despite formulations to optimize cell relationships and tailored regularizations of those previously proposed^16^ these are no implementations based on DNA sequence inputs. No DL methods in single-cell genomics jointly optimize DNA sequences and cell relationships, due to scaling limitations, and it is needed to computationally decode regulatory rules that guide cellular states^17–19^.

To overcome these limitations, we introduce *muBind*, a single-cell compatible method designed to predict genomics counts from sequence features and graph representations. Inspired by the original *ProBound*, a core innovation of *muBind* is the introduction of a *Graph Layer* that optimizes and propagates cell-specific sequence activities across nearest-neighbor using sparse tensors. Benchmarks against *PyProBound* in bulk genomics and *scBasset* in single-cell genomics, demonstrate the competitive performance of *muBind* in multiple dataset setups. Leveraging cell dynamics priors derived from RNA-velocity shows that *muBind* can identify motif-graph regulatory features, interpreted as dynamics-associated TFs supported by orthogonal evidence in biological case studies, like Sox9 role in early pancreatic endocrinogenesis, and Gli3/Prdm16 in neurogenesis and human organoids. *muBind* is a robust method for sequence-to-activity inference in single-cell genomics, improving the understanding of cell dynamics.

## Results

### A scalable method for DNA motif inference in single-cell genomics

We developed *muBind*, a PyTorch-based model inspired by *ProBound*, to process single-cell genomics data using interpretable DNA sequence-by-cell representations and *k*-nearest neighbors (kNN^20,21^) graphs (**Fig 1a**). *MuBind* implements a regression task to predict genomics counts from input sequences (**Fig 1b**), using three general building blocks. The first two are akin to *Probound*: (i) a *Binding Layer*, where mononucleotide and/or dinucleotide sequence representations pass through a series of independent convolutional filters, with random or pre-declared, tunable weights, (ii) an *Activities Layer*, where learned filters are weighted by their relative contribution per sample. Importantly, the primary innovation of *muBind* is (iii) a *Graph Layer*, which propagates sequence-based activities (*Y*) to neighboring cells using kNN-graphs. This new operation is accomplished via matrix multiplication of the Activities Layer output by the Hadamard product of a prior connectivity graph (*G*) and a learnable graph tensor (*D*), both represented using sparse tensors (**Methods**), allowing light-weight computing of single-cell genomics graphs, with desirable upper memory boundaries. *MuBind* supports learning motifs *de novo* or refining known position-weight matrix databases from ENCODE^22^ (**Methods**). It implements greedy optimization in the Binding Layer via iterative sub-groupings, weights freezing, and reshaping. By adopting scverse^23^ guidelines, *muBind* processes single-cell genomics data using *bedtools* and *AnnData*, offering modular, scalable single-cell modeling and analysis.

**Figure 1.**
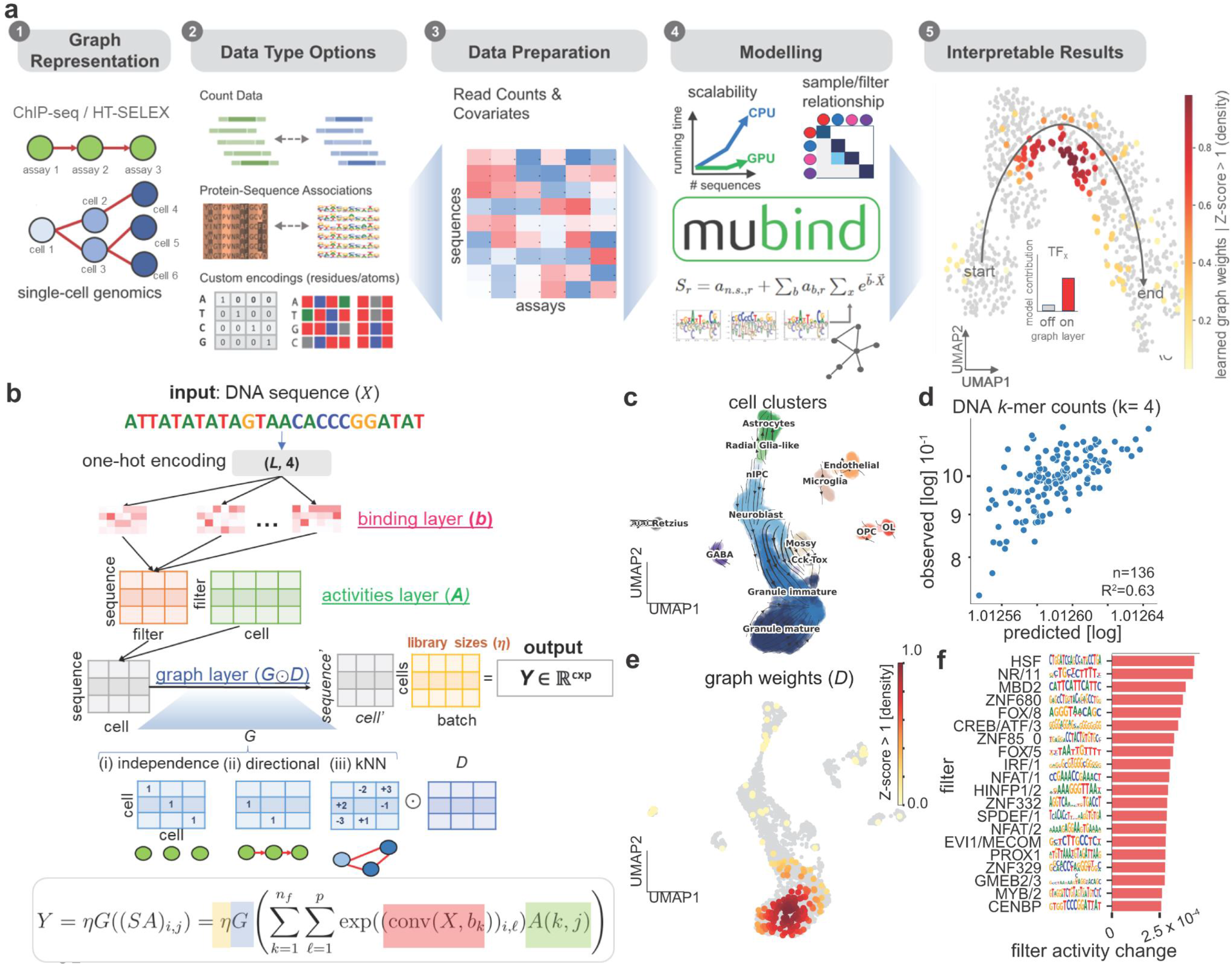
*muBind* is a method for sequence-to-activity modeling using graph representations in single-cell genomics. (**a**) (1) Representation of genomics samples (bulk assays or single-cells, nodes) and their relationships (edges). In bulk genomics (green nodes, ChIP-seq, HT-SELEX, and similar) relationships between assays are explicit (red arrows), whereas in single-cell genomics (blue nodes) connectivities are inferred and stored as *k*-nearest neighbors^20^ (red lines) (2) Data ingestion and representation of genomics data is majorly based on DNA reads and count tables, motivating (3) sequence-by-cell representations as input for modeling methods. (4) *muBind* is a regression model that can learn motif contributions from such input representations, via learnings of motifs and graph-based sample contributions. (5) Output interpretability. *muBind’s* learned graph layer weights can be inspected per cell (UMAP, color bar) and by comparison of net filter activity changes upon activation (inset barplot). (**b**) *muBind* model architecture. Visualization of input DNA sequences (X) and its way through the model to generate outputs *Y^*(cxp) (cell times peaks). One-hot encoded DNA sequences pass through three main building architecture blocks: The *Binding* layer (*b*, salmon), the Activities Layer (*A*, green) the *Graph Layer* (*G*, blue), and a library size correction (*η*, orange). The bottom formula summarizes the core operations of each layer in matched colors. *G* main parameters (tensors G and *D*) are shown as matrices but are implemented via sparse representations. Additional equations, regularization terms, and optimization details are further described in **Methods**. (**c**) dentate scRNA-seq gyrus dataset visualized with vector stream obtained from RNA-velocity graph. (**d**) *muBind* predictions (x-axis) of observed counts (y-axis) for all DNA *4*-mers (n=136) listed in the dataset. (**e**) Visualization of cells with high absolute sums of learned graph weights (*D*), shows as Z-scores (Z > 1). (**f**) normalized activity changes of binding layer filters of *muBind* models trained with versus without a Graph Layer.

Using mouse dentate gyrus as an example^24^ (**Fig 1c**), *muBind* predicts observed counts gene using DNA sequences from TSS-associated regions (**Fig 1d**). Observing the weights learned in the Graph Layer *D* allows to annotate cell clusters with high contributions and potential kNN-based propagation of sequence activities (**Fig 1e**). Finally, activity changes per filter versus a model trained without a Graph Layer (**Fig 1e**), or via weighted activity-graph contributions (**Methods**), can be used to relate binding filters with the learned graph weights.

### Benchmarking *muBind* in bulk genomics

To demonstrate the robustness of *muBind*, we benchmarked it against state-of-the-art methods for bulk (*ProBound*) and single-cell (*scBasset*) DNA sequence-based prediction of read counts. When comparing *muBind* and PyProBound *models*, we observed high agreement in R^2^ values for HT-SELEX count predictions across datasets (n=100, R=0.81, *P*<0.001) (**Methods, Supp Fig 1a**), with learned motifs and activities from *muBind* aligning with the underlying TF-binding specificity of TFs profiled (**Supp Fig 1b-d**). These results indicate that *muBind* with its *Graph Layer* can reproduce the core prediction capabilities of *ProBound* in bulk genomics, encouraging its application to single-cell datasets with complex kNN-graph representations.

### Benchmarking *muBind* in single-cell genomics

After confirming the reliability of *muBind* in bulk genomics, we benchmarked its performance and Graph Layer scalability in single-cell genomics. We prepared four scATAC-seq datasets based on droplet-based sequencing for demonstrating applicability: (i) PBMC (10,246 cells^25^), (ii) mouse neuronal differentiation (5,877 cells^26^), (iii) mouse pancreatic endocrinogenesis (16,918 cells^27^), and (iv) neuronal organoids (34,088 cells^17^), formatted as sequence-by-cell AnnData representations, based on *sfaira*^*28*^ principles. For comparisons, we selected features using two approaches (*random*, or *episcanpy*^*29*^) and trained *muBind* models with two hyperparameter combinations: Binding layer (*de novo* or *pwms*), and activation of the Graph Layer with a pre-trained nearest-neighbors kNN-graph (*graph*). For comparisons, we selected *scBasset*^*10*^, which classifies accessibility levels from DNA sequence features using a convolutional tower followed by a final embedding layer. As *muBind* uses a Poisson loss to learn observed counts^30^ in addition to testing *scBasset* using its default binary-cross entropy loss (*bce*), we also tested using Poisson (*poisson*) (**Methods**).

In 80/20 training/testing splits (five-fold), muBind outperformed scBasset in prediction performance (R^2^) and overall score across all datasets (**Supp Fig 2-3**), consistent with quantitative losses better modeling genomics counts^30^. *scBasset* with Poisson loss did not yield better results, suggesting overfitting or additional hyperparameter tuning is needed. The classification performance of *muBind* also outperformed *scBasset*, with greater PR-AUC values in most cases. The *Graph Layer* might decrease generalization performance due to additional parameters, yet it negatively affects the performance of the best model configuration (*pwms+random*) only in one dataset (Organoids, 0.41 versus 0.48, respectively), while overall scores are improved or maintained in others (0.26 versus 0.25, one case; 0.30, two cases)), indicating minimal generalization issues and competitive running times of *muBind* (**Supp Fig 5**).

**Figure 2.**
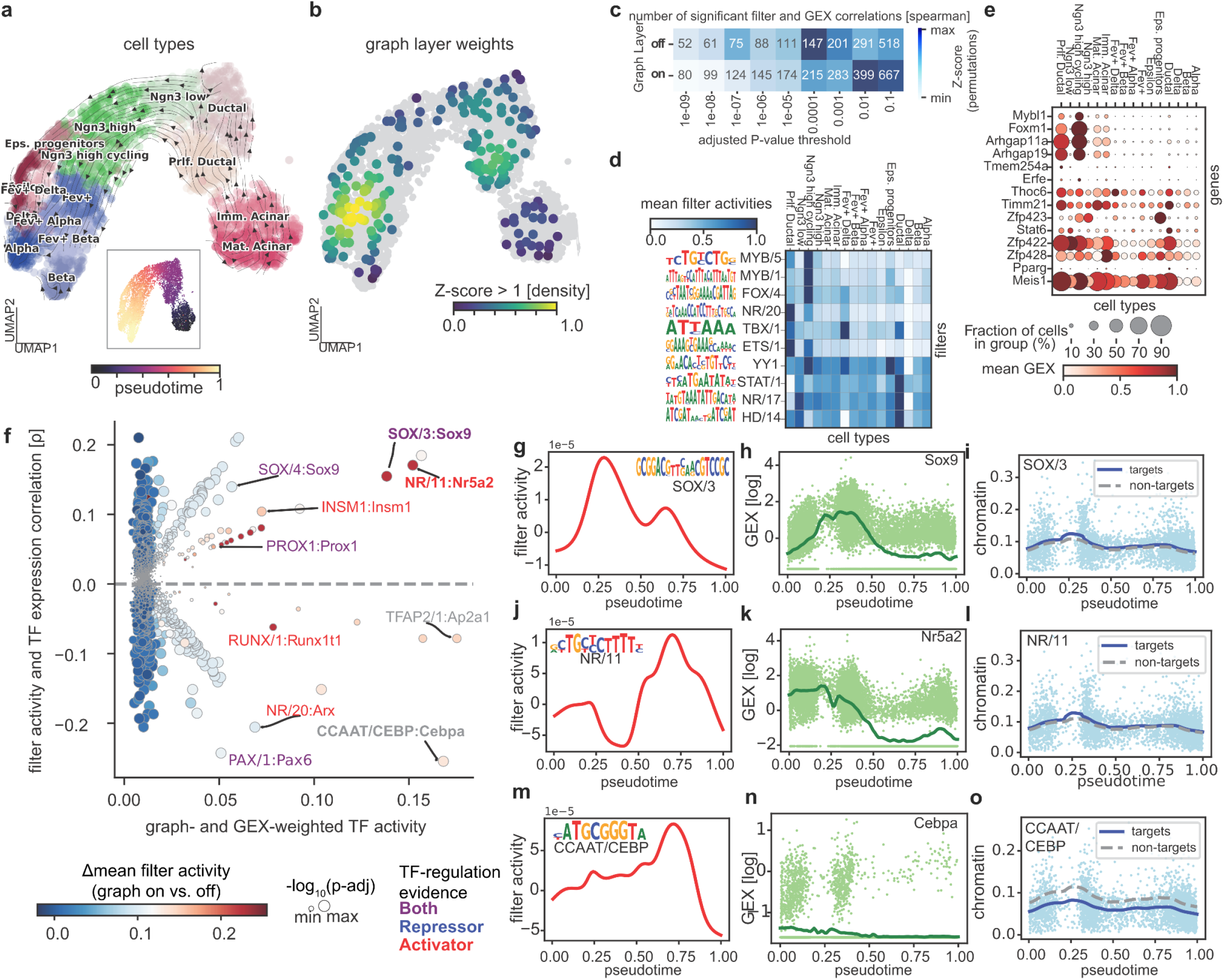
*muBind* learns dynamics-weighted sequence features that predict chromatin activity in pancreatic endocrinogenesis. (**a**) UMAP visualization of the pancreatic endocrinogenesis dataset with velocity streams calculated from RNA (*scVelo*). (**b**) Absolute values of graph layer learned weights (*D* tensor, see **Methods**), visualized a sum of absolute weights per cell (Z-scored) and as a density using a fixed threshold (Z > 1) (**c**) Number of significant *Spearman* correlation pairs between filter activities and matched TF gene names, using gene expression values, for models with and without a Graph Layer (*y*-axis), across different *P*-value thresholds (*x*-axis). *Z*-scores highlight value differences versus an empirical correlation distribution against decoy TFs from non-related families. (**d**) Mean filter values are averaged per cell type. Filter labels are based on ENCODE^37^ motif archetypes. Logos indicate refined binding layer weights. (**e**) Gene expression of matched TFs from top-enriched filters from *c* with high value-value correlations. (**f**) *Y*-axis: Spearman correlation between gene expression and filter activities. *X*-axis: Combined product of filter-graph interaction weights (**Methods**), the absolute sum of per-cell filter activities, and correlation between gene expression and filter activity (cell-wise). Color scale: Mean change in filter activity weights for models with a graph layer versus without. Circle sizes: Significance of the correlation between filter activities and gene expression. Text labels: modes of TF gene regulation (sourced from GRNPedia^38^). Text labels show filter:TF cases with high-combined scores (x-axis), and high effect correlations between filter activity and gene expression (color bar) (**g**) Mean filter activity of the SOX/3 module against pseudotime. (**h**) Gene expression of the Sox9 gene over pseudotime. Each dot indicates one cell. (**i**) Chromatin activity of SOX/3 targets versus non-targets, selected using high-quantile binding scores. (**j**) Similar to *g*, showing the NR/11 module. (**k**) Similar to (**h**) showing Nr5a2. (**l**) Similar to *i*, showing NR/11. (**m**) Like *g*, showing the CAAT/CEBP module. (**n**) Similar to *h* showing Cebpa. (**o**) Similar to *i*, showing CAAAT/CEBP.

### *MuBind* and RNA-velocity reveal sequence-to-dynamics associations in single-cell genomics

*MuBind* learns interpretable motif filters (as *ProBound*, **Supp Fig 1**) and is competitive versus *scBasset* in single-cell genomics, without major generalization penalties from the Graph Layer (**Supp Fig 2-3**). To test the effectiveness of the Graph Layer, we trained models to learn the propagation of sequence activities through biologically close cells using kNN-graphs and the learnable matrix *D* from the graph layer, without additional constraints. This analysis did not show directionality and coherence in velocity vectors derived from the Graph Layer, suggesting that additional, pre-defined pseudo-time-based representations can be useful while learning DNA sequence contributions to cell dynamics (**Supp Fig 4e,k**). This is consistent with chromatin accessibility work that shows scATAC-seq dynamical modeling is improved once additional states of transposase chemical reactions are included, or alternative modeling parameters are considered^31,32^, and motivated us to complement *muBind* models with scRNA-seq dynamical features.

We trained *muBind* models using a Graph Layer where kNN-connectivities are replaced with RNA dynamics-based graph representations from *scVelo*^*33*^ (velocity graph). This updates *muBind’s* Graph Layer *G* matrix with a dynamics prior (**Methods**) and detects sequence-to-activity contributions across neighboring cells via the learnable matrix *D*. As *muBind* can learn models from sequence-by-cell representations, we tested the addition of the velocity-graph on datasets in two organisms and modalities: (i) mouse pancreatic endocrinogenesis with scATAC-seq (**Fig 2**), and (ii) neurogenesis with scRNA-seq counts, in mouse and human organoids (**Fig 3**). In both cases, we tested the Graph Layer usability by interpreting interactions between the filters in the binding layer with graph weights, relationships with matched TFs and their gene expression, and their dynamics over pseudotime.

**Figure 3.**
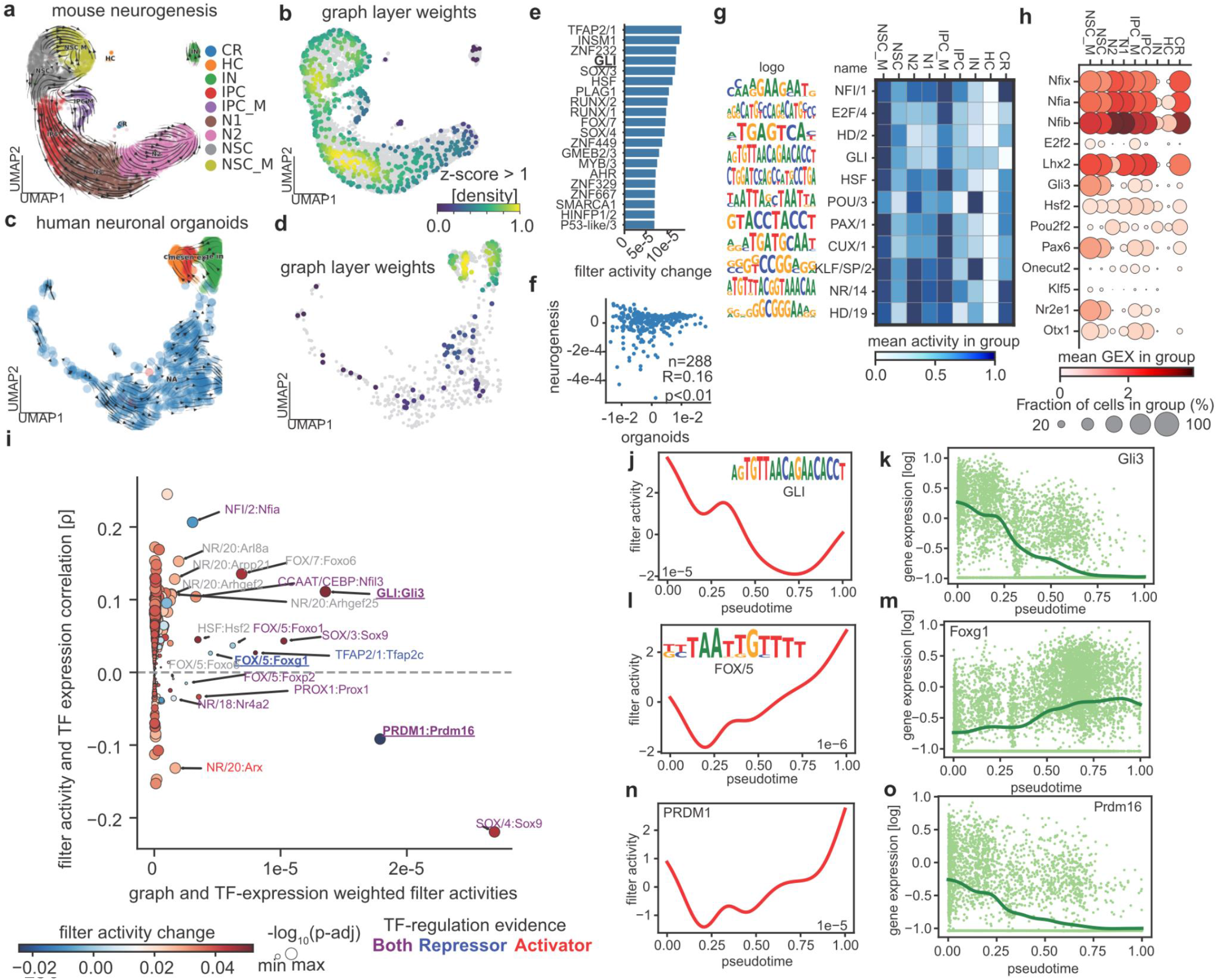
*muBind* learns species-reproducible motif-graph relationships that predict cell transitions in neurogenesis scRNA-seq. (**a**) UMAP of neurogenesis^26^ showing *scVelo* velocity vectors. CRs, Cajal-Retzius neurons; MG, microglia; IPC, Intermediate Progenitor Cells; NSC, Neural Stem Cell. N, Neurons. _M indicates mitotic sub-groups. (**b**) Graph Layer weights are summarized as absolute values on the embedding. (**c**) UMAP of human neuronal organoids^17^ showing velocity embedding (*scVelo*) (**d**) Graph Layer weights learned by *muBind*. (**e**) Mean filter activity difference between muBind models with an activated Graph Layer, versus models without one (see **Methods**). Labels are based on ENCODE motif archetypes. (**f**) correlation between graph layer contributions to filter activity changes in mouse neurogenesis (y-axis) versus human organoids (x-axis). (**g**) Mouse neurogenesis mean filter activities are summarized per cell type. Visualization of motifs depicts logos of mononucleotide-refined motif weights. (**h**) Gene expression of filter-matched TFs that correlate with filter activities, calculated using per-cell readouts. (**i**) Selection of genes whose filter activities are associated with high per-cell filter activities, filter weights, and correlation with TF gene expression. Legends as label annotations as in **Fig 2f**. (**j**) Filter activity of the GLI motif archetype. The line indicates the mean across cells. (**k**) Gene expression of Gli3 gene over pseudotime. Each dot indicates one cell. (**l**) Filter activity of the FOX/5 module. The line indicates the mean across cells. Gene expression of the Foxg1 over pseudotime. Each dot indicates one cell. (**n**) same as *j,l* with PRDM1. (**o**) Same as *k,m* with Prdm16.

### Pancreatic endocrinogenesis

We asked if *muBind* using a Graph Layer pre-loaded with an RNA-velocity graph can identify TFs driving pancreatic endocrinogenesis chromatin dynamics (**Fig 2a**). Performances were comparable with and without the *Graph Layer* (**Supp Fig 4d**). We summarized *Graph Layer* learned weights from *D* in three ways: (i) absolute sum of weights *D* per cell, normalized as Z-scores (*graph weights*) (**Fig 2b**), (ii) associations between motif filter activities and graph activities per cell (*filter-graph contribution score*) (**Fig 2f, Methods**), and (iii) binding layer weights with graph weights and gene expression for matched TFs (**Fig 2d-e**).

Cell with high graph weights clustered in two lineage-branching sectors (proliferative ductal to Ngn3 low, and Ngn3 high-cycling to Delta/Alpha/Beta), indicating the Graph Layer captures relevant dataset regions that could be enhanced with neighboring sequence-contributions during cell transitions (**Fig 2b**). Gene expression validation showed stronger associations between binding layer filter activities and matched TFs with the Graph Layer when compared to an empirical distribution of decoy TF genes (**Fig 2c**), suggesting that regulatory factors linked with cell states can be more specifically detected (**Fig 2d-e**). Combining the cell-wise graph contribution and filter TF activities allows us to focus on candidate regulators whose activities are linked with Graph Layer weights (**Supp Fig 5**). This, multiplied with the absolute sum of filter activities, and TF gene-expression concordance with filter activities, allowed us to calculate a single combined product of the three metrics, for visualization and prioritization of TFs with high relevance in explaining graph-based predictions (**Fig 2f**). This scoring revealed among top candidates Sox9, a known early regulator in ductal cells that induces Ngn9, as one of the main graph-enhanced regulatory factors of the cell transition^34^. This factor has been studied in mouse knockout models due to genetic associations between human SOX9 mutations and pancreas developmental disorders^35^, however, this insight has not been observed before via single-cell RNA-seq dynamical analysis, suggesting the Graph Layer enhances biological discovery via sequence-based modeling of cell dynamics. We validated this association further by examining the genome regulation across pseudotime of Sox9 by visualizing SOX/3 filter activities, Sox9 gene expression, and chromatin accessibility of SOX/3 target loci (**Fig 2g-i**), revealing a high concordance of these three values at earlier pseudo-timepoints. Other TFs like Nr5a2 (**Fig 2j-l**), known for its role in exocrine development^36^, and negative gene expression and chromatin associations with Cebpa (**Fig 2m-o**) were also detected (filters NR/11 and CCAATT/CEBP, respectively), underscoring the ability of *muBind* to prioritize sequence feature effects on cell transitions with high specificity.

### Neurogenesis

We explored the ability of *muBind* to discover TFs driving neuro-cortical cell development from neuronal stem cells, using mouse neurogenesis (E14.5)^26^ and human neuronal organoids^17^, where RNA and an RNA-derived velocity graph are available. (**Fig 3a,c**). As a proxy for genomic regulation, *muBind* can be trained using Transcription Start Site (TSS)-associated DNA sequences of mapped genes. This analysis revealed graph weights related to regions of suspected cell transitions, like NSCs towards IPCs and late organoid stages (**Fig 3b**). The activation of the graph layer highlighted motif filters that contributed to cell-type-specific regulatory changes (**Fig 3e**) and associated those with regulatory changes in cell-type-specific gene expression levels of filter-matched TFs (**Fig 3g-h**). Importantly, *muBind* models for scRNA-seq human organoids showed a significant correlation for graph-associated filter weight activities with mouse neurogenesis, and top candidates belonging to the same TF families in both datasets (R=0.16, *P*<0.01). This suggests that *muBind* can reconcile similar motif-graph interactions across species (human vs. mouse) and conditions (*in vivo* vs. *in vitro*) (**Fig 3f**).

Global binding filter activities were linked to mouse neurogenesis TFs (**Fig 3g,h**). Similar to pancreas endocrinogenesis (**Fig 2f**), by jointly assessing filter-graph weight interactions (**Methods**), filter activities, and filter gene-expression associations, we found candidate TFs associated with regulatory control of RNA dynamics (**Fig 3i**).

Specifically, Gli3 is a known early neurogenesis activator^39^, also tested in organoid-specific perturbation experiments^17^. This prediction was further reinforced by the association between GLI filter activities and Gli3 gene expression over pseudotime (**Fig 3j-k**). Multiple members of the Forkhead family are also detected among main graph-regulatory candidates. Among these, Foxg1 has a strong link with adult neurogenesis maturation^40^, whereas Foxo1 is related to memory and synaptic activity but is not crucial during neurogenesis ^41^. Expression data of Foxg1 over pseudotime confirms a correlation with filter activities at the late stages of the differentiation, highlighting its role as a non-redundant regulator of cortical development^42^ (**Fig 3l-m**). Finally, among repressor TFs *muBind* detects PRDM1-related motifs contributing to graph-based model predictions, highlighting Prdm16 as a candidate dynamical regulator. The high expression of Prdm16 at early pseudo time points concomitant with increasing PRDM1 filter activities further highlighting this TF as an early genomic regulator of early neurogenesis^43–45^. Filter activities increasing at late pseudotime points can be interpreted as PRDM1-induced gene programs remaining active long after Prdm16 induction and sequential down-regulation.

Overall, incorporating the *Graph Layer* in *muBind* enhances its informativeness and scalability for single-cell genomics, effectively prioritizing sequence-specific, dynamics-associated TF regulation during cell transitions.

## Discussion

We introduce *muBind*, a model inspired by a biophysically interpretable machine learning algorithm^12^ now enabling single-cell genomics analyses with kNN-graph representations. The *Graph Layer* is a major upgrade, facilitating joint learning of relationships between samples and sequence features, particularly useful for understanding cell transitions. We provide repositories for data loading, benchmarking, and tutorials for *muBind* (see **Code Availability**). *muBind* analyzes genomics assays of any complexity, using sequences and sample matrix representations. It applies to scRNA-seq by transforming gene features with gene promoter sequences, as demonstrated in neurogenesis. Comparisons with *Probound* using HT-SELEX datasets indicate that other bulk genomics modalities, such as ATAC-seq and ChIP-seq, are also suitable, offering additional promising avenues in joint modeling of bulk and single-cell epigenomics.

The Graph Layer in *muBind* supports sparse tensor graph representations, allowing the declaration of *de novo*, kNN-based, or RNA-dynamics-based weights to learn sequence-to-activity associations. While *ProBound* used sequential weight propagation for bulk experiments, kNN-graphs are most used in single-cell genomics and require additional memory, (*n*^*2*^ with the number of cells, in the worst case). Our Graph Layer implementation, utilizing sparsity operations, can handle current single-cell genomics scales (between 10^5^-10^6^ cells and 10^4^-10^5^ features). Future work must explore improvements to our current graph regularization approach and other graph properties to further compute and assess the significance of graph-sample and graph-motif predictions.

Chromatin accessibility readouts alone showed limitations in learning graph weights that can used to interpret cell transitions, aligning with chromatin-only dynamics being less informative than RNA-based readouts for detecting biological directionality until custom protocols^31^ and model modifications are tested^32^. RNA-based velocity weights in the Graph Layer of *muBind* helped improve sequence-to-activity predictions, allowing the global classification of cells into dynamic or static categories by inspection of the absolute sum of learned graph weights per cell (**Fig 2b** and **3b,d**). We envision CellRank2-derived kernels as graph priors in the *Graph Layer* to train *muBind* models and infer associations between sequence features and readouts in cell transitions^15^.

Previous work has proposed learning dynamical features using a prior TF-gene regulatory network representations^18,19,46^. On this aspect, one key distinction of *muBind* is that it can simultaneously optimize DNA binding weights and activities in one modality (scATAC), from nearest-neighbors weights, and/or priors from other layers (RNA-velocity). Using a custom scoring of filter-graph interactions, we reconciled cell-specific and dynamics-associated TFs, highlighting key transcriptional regulators like Sox9/Nr4a2 in pancreatic endocrinogenesis, and Gli3/Foxg1/Prdm16 in neurogenesis. Dynamics-independent TFs may result from chromatin-independent co-regulatory effects that maintain cell-identity without being crucial for process directionality, or correlation artifacts due to scRNA-seq detection limits of transcription factors.

One limitation of *muBind* is distinguishing biologically meaningful filters from sequencing biases (e.g. GC-content). Due to multiple interacting TFs, this regression task fits *de novo* motifs and activities in HT-SELEX more often than in single-cell genomics. Future work should include optimization guidelines and benchmarks with more complex regularizations, like dropout-based continuous sparsity^47^, and active model refinement methods. Another limitation of *muBind* is the lack of batch correction for integrating biologically equivalent samples from different experiments. While the last layer of *muBind* includes batch information as a matrix, its performance is limited, suggesting complementing with variational autoencoders approaches as further work (e.g. scVI^48^, PoissonVAE^30^). Benchmarking biological sequence representations and their role in batch correction has not been fully evaluated compared to scRNA-seq^49^ using genes as tokens. We foresee developing unified sequence-based representations and identifying potential sequence-based mutual nearest neighbors^50^ in single-cell transitions.

A final, exciting aspect of our work is the potential for full multiome-aware models fits with *muBind*. Using concatenations of ATAC and RNA cells as sequence representations is possible, yet current dataset availability has limited our testing on combined sequence feature spaces with interacting factors. Extensions of the Activites Layer, provided as tutorials in the current release, allow incorporating additional modalities in joint-modeling tasks. Further exploration of interactions terms, losses, and batch correction regimes across multiple biological systems with paired multiome data will confirm the usability and robustness of *muBind* in more complex setups, paving the way for understanding regulatory genomics from single cells.

## Methods

### Model details of *muBind*

*muBind* implements a sequence-to-activity model, where DNA sequence features are encoded as one-hot features. Input *X* ∈ ℝ^*s*×*p*^ with *s* representing sequences and *p* positions in those are used to predict a matrix of cells *Y* ∈ ℝ^*c*×*p*^ where *c* is cells (or assays). This is done through a series of three neural net layers:

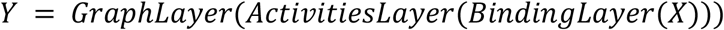

### Binding and Activities layer

The binding layer applies a series of parallel, independent kernel operations on *X*, using pre-declared, trainable weight matrices 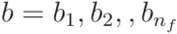, where ^*n*^_*f*_ defines the number of kernels. The output of these independent operations is stored as a tensor 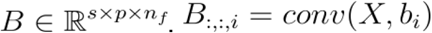 with *i* ∈{1,,*n*_*f*_}, where conv() denotes the convolution.

Kernels in b have dimensions (w, m+d), where 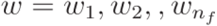, and indicate the number of mononucleotide (4) or dinucleotide features (16) per position, respectively. Thus, each kernel can require between 4w and 16w independent weights per position, depending on whether dinucleotide encodings are activated within each layer.

To compute the overall binding score for each kernel, and akin to *ProBound* we compute element-wise *exp*(*B*) and sum over the positions in the sequence, such that we receive one score per sequence and kernel. The result is stored as a matrix 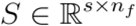:

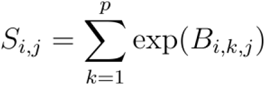

The implementation of this layer utilizes the *Conv1D* module function in PyTorch. The declaration, extension, and transformations of weights in this layer are iteratively optimized through weight freezing and refinements, following sequential strategies described by *ProBound*. In this work, we always optimized one filter at the time in the *de novo* setting, including shifting and width extensions. When filters are preloaded from a motif library (*pwms*), these are optimized in groups of 20 filters at the time.

The activities layer defines a multiplication of S by a matrix 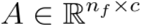, converting S into a matrix of *Y*_*sample*_ ∈ ℝ^*s*×*c*^. This matrix captures the relative contribution of each binding filter in each cell for cell-based predictions.

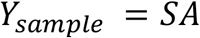

### Graph Layer

The graph layer performs operations to capture sample-sample relatedness, leveraging a predefined graph. Two tensors are defined in this layer (i) a graph matrix *G* ∈ ℝ^*c*×*c*^ that specifies the true connections between samples, and (ii) a learnable weight matrix *G* ∈ ℝ^*c*×*c*^, which is used to declare the relevance of these connections for prediction. The main operation is defined as the product of *Y*_*sample*_ and the Hadamard product of *G* and *D*.

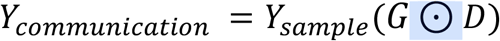

Conceptually, the diagonal of *G* captures self-sample memberships, while off-diagonal values handle sample-based relationships in both directions. As additional and optional constraints in *D*: (i) entries *D*_*i,j*_ are only necessary when *G*_*i,j*_ values are not zero, and (ii) *D*_*i,j*_ = − *D*_*j,i*_. These two constraints significantly reduce the number of parameters versus a dense matrix representation.

The Graph Layer is implemented as `tnn.Parameter` () values, which are reshaped into sparse matrices, and multiplied accordingly using `torch.sparse` modules. This approach reduces the total memory footprint required during training operations compared to the dense matrix approach. When tested on a portable NVIDIA card with 16 GB of memory and assuming a batch size of 512 or lower, we can train our largest dataset examples, organoids scATAC-seq (340k cells and 550K peaks). The total estimated time, based on benchmarks, is approximately six times the training time of a *muBind* model without a Graph Layer (**Supp Fig 5**).

Altogether, the three main operations of *muBind* can be summarized as a single term, including a learnable vector *η* ∈ ℝ^*c* ×1^ to adjust predicted values for library sizes

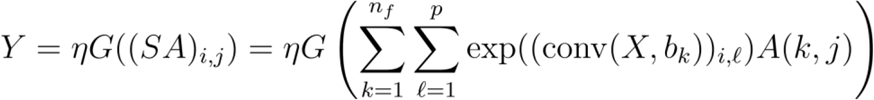

### Loss function

The combined loss function for *muBind* is the sum of multiple individual losses that can be deactivated based on user-specific configurations, and can be calculated in parallel. Among the most relevant ones, we highlight the following four:

- prediction error between observed and predicted read counts, using Poisson Loss.
- minimization of overall weights similarities in the binding layer (filter dissimilarity).
- graph layer regularization: this term accounts for differences in values D, conditioned by valid nearest-neighbor interactions between sample pairs (i, j) and any other pair (i, k). This is calculated using the following sum:

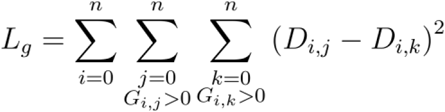
- Dirichlet and/or exponential barrier constraints per column for weights in binding layer filters

During training, after each epoch, losses are combined using a joint sum 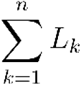. In this work, the three main losses are prediction error, graph regularization, and filter dissimilarity (**Supplementary Figure 2**), with an exponential barrier constraint. Combinatorial testing of additional loss setups is possible, and we refer to additional running settings in the current release documentation.

### Interpretation of activity and graph layer weights in *muBind*

#### Binding layer and Graph Layer synchrony

To understand how each filter contributes to the overall prediction, we note that by definition

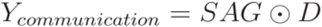

For a set of activities linked with a filter *a*_*i*_, i.e. a row of *A*, to have a meaningful impact on *Y*_*communication*_, it needs to:

1. be well aligned with the weighted Graph Matrix *G* ⊙*D*: For the i-th row of *A*, denoted as *a*_*i*_, the quotient 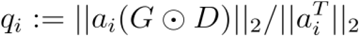 should be relatively large.

2. Have a large 𝓁^1^ norm compared to other filters. 𝓁^1^ norm compared to other filters.

Since *A*(*G*⊙*D*)=((*G*⊙*D*)^*T*^*A*^*T*^)^*T*^, we compare *q*_*i*_ to the maximum singular value σ_*max*_ of (*G*⊙*D*)^*T*^., to ensure values between 0 and 1. Finally, we compute the product 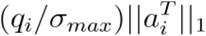 to find filters that fulfill both properties. *A*(*G*⊙*D*)=((*G*⊙*D*)^*T*^*A*^*T*^)^*T*^, we compare *q*_*i*_ to the maximum singular value *σ*_*max*_ of (*G*⊙*D*)^*T*^, to ensure values between 0 and 1. Finally, we compute the product 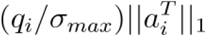 to find filters that fulfill both properties.

The calculation of these contributions is implemented in the library method `mubind.tl.compute_contributions`

### Graph Layer scores distributions in dataset

Graph *D* weights are interpreted within a dataset by assessing the distribution of high absolute values through kNNs. This is visualized using the method `embedding_density` in Scanpy^20^.

### Binding activity changes upon Graph Layer activation

To write the column-wise difference between the scores in two activity matrices, we denote the two matrices as (*A*) and (*B*), each with dimensions (*p* ×*nf*). Briefly,

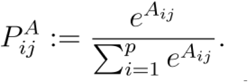

Let (*A,B* ∈ ℝ^*p*×*nf*^) be the matrices.

-Let 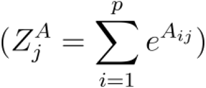 and 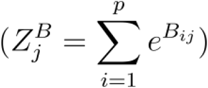 be the partition functions for column \(j \).

Let 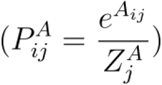 and 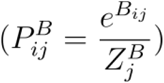 be the normalized scores.

-The column-wise difference (*D*_*j*_) is then given by:

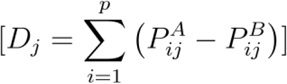

In mouse neurogenesis, the mean of the final value changes per filter is visualized on **Fig 2e**.

### Graph- and gene-expression-weighted TF-activity

Using (i) graph and binding layer syncrony per filter, (ii) the global correlation between gene expression and binding filter activity per filter (Spearman correlation), and (iii) the mean change in absolute filter activity between a model without and with the Graph layer, we use the combined product of these three values as a score to prioritize and interpret TFs. This is shown in the x-axes of **Fig 2f** and **3i**.

### Comparison of *muBind* and *ProBound*

Using datasets and models from the MotifCental database (motifcentral.org) as a reference, we listed all 582 HT-SELEX^51^ models and retrieved FASTQ files linked with those. Using repositories for data retrieval (*bindome*) and benchmarking (*mubind-benchmark*), we tested *muBind* on 100 random HT-SELEX datasets from those, using either 10,000 (this work) and up to 100,000 unique sequences per dataset (*mubind-benchmark*). In both *muBind* and *PyProBound*, we tested various hyperparameters before selecting the model with the higher R^2^: (i) number of binding layer filters (1 to 5), (ii) dinucleotide encodings (on/off), (iii) filter width optimization, and (iv) filter weight shift optimization.

### Comparison of *muBind* and *scBasset* in single-cell genomics data

Four single-cell chromatin accessibility datasets were retrieved and reprocessed for comparison between *muBind* and *scBasset*: (i) PBMC (10,246 cells^25^), (ii) mouse neuronal differentiation (5,877 cells^26^), (iii) mouse pancreatic endocrinogenesis (16,918 cells^27^), and (iv) neuronal organoids (34,088 ATAC metacells^17^.). During preprocessing, dataset samples were prepared with combinations of the three following parameters: (i) number of cells = {100, 500, 1000, 5000, 10000, and all}, (ii) number of ATAC-seq peaks (three times the number of cells), and (iii) selection rule of highly variable regions {episcanpy^29^, random}. Results of the benchmark shown in this work are based on 5,000 cells and 15,000 features, and further tables are deposited in the benchmarking repository. We used the GitHub release of scBasset (commit ID: d31138b). Training, validation, and testing groups were prepared using the *scBasset* script (scBasset.py). Custom incorporation of a Poisson loss into *scBasset* was done using PoissonLoss as implemented in PyTorch. Training was done using CPUs. To compare *scBasset* and *muBind*, we used classification metrics (ROC-AUC, PR-AUC, multi–label) and quantitative features (R^2^). These values are calculated on the training and testing fractions, prepared using *scBasset’s* default 80/20 train/test splits.

Testing the Graph Layer is possible with *muBind* because training/testing splits are prepared using sequences, and not samples. In benchmark cases, the kNN-graph was prepared using Scanpy^20^ with default readouts.

### Incorporation of external graph information

To test the incorporation of a custom velocity graphs in *muBind’s* core model, we processed a multiome scRNA+ATAC^27^ pancreatic endocrinogenesis dataset using *scVelo*. A velocity graph *V* ∈ ℝ^*c*×*c*^ updates the weights of G in the Graph Layer, allowing the weights *D* to be tuned during the learning process in *muBind*. In the case of Neurogenesis and organoids, the graphs are available in the respective publications^17,26^.

### Interpretation of motif-graph interactions

Filters in the *muBind* binding layer can be learned *de novo* or pre-loaded and refined using a library of position weight matrices. In this work, we use and provide loaders to include ENCODE motif archetypes^22^ (n=288). When using this pre-loaded library, filters and activities are directly correlated with the gene expression of groups of transcription factors.

We compare the activity of each binding filter with the gene expression of corresponding transcription factors by calculating per-cell RNA and filter activity correlations, as well as chromatin accessibility and filter activity correlations. To calculate the significance of this, we simulate 20 times decoy relationships between filters and non-matched TFs, and recalculate the number of observations. Association between TFs and gene regulation mode (activator or repressor) is retrieved and shown based on annotations from GRNPedia^38^.

To assess the influence of filters across pseudotime, we measure chromatin accessibility of targets vs non-targets for a given binding filter, using at least the top-20 quantile of binding scores across all ATAC-seq peaks for that filter.

## Code and Data Availability

*muBind* is implemented in Python as a PyTorch model, and it is publicly accessible at www.github.com/theislab/mubind, Tutorials for bulk, single-cell genomics data are available at www.readthedocs.io/mubind.

The reproducibility of bulk genomics benchmark results presented in this work is available through a Snakemake workflow, accessible at https://github.com/theislab/mubind-benchmark. Independent Jupyter notebooks are also available for data preparation and benchmarking of single-cell genomics datasets discussed in this work.

Data retrieval and manipulation of genomics and sequence data for machine learning purposes can be done using bindome, available at www.github.com/theislab/bindome, with tutorials at www.readthedocs.io/bindome.

## Acknowledgments

We thank members of Fabian Theis’s lab for their constructive feedback on this project. We thank all the beta testers of *muBind*. We thank Natalie Romanov, Mariia Minaeva, Anastasia Litinetskaya, Isaac Virshup, Soroor Hedizeh-Zadeh, and Fabiola Curion for suggestions for manuscript input and package improvement suggestions. We thank Faye Chong and Jonas Fleck for their valuable input while preparing the mouse neurogenesis and human organoids scRNA-seq datasets. I.L.I. acknowledges the funding and support provided by Wellcome Leap (Delta Tissue).

## Author Contributions

I.L.I. conceived the project and led all execution aspects. J.S. assisted in implementing the prototype version of *muBind*. E.E developed the first snakemake workflow for comparing *muBind* and *ProBound*. L.M. assisted in the interpretation of results from *muBind* and *scBasset* using Poisson models. H.A. contributed to the theoretical conceptualization and testing of dropout-based regularizations in *muBind*. D.K. prepared the pancreatic endocrinogenesis dataset and assisted in the interpretation of results. L.H. conceptualized and implemented graph-filter contribution metrics. F.J.T contributed with funding, mentorship, and fostering lab synergies.

## Supplementary Material

**Supplementary Figure 1.**
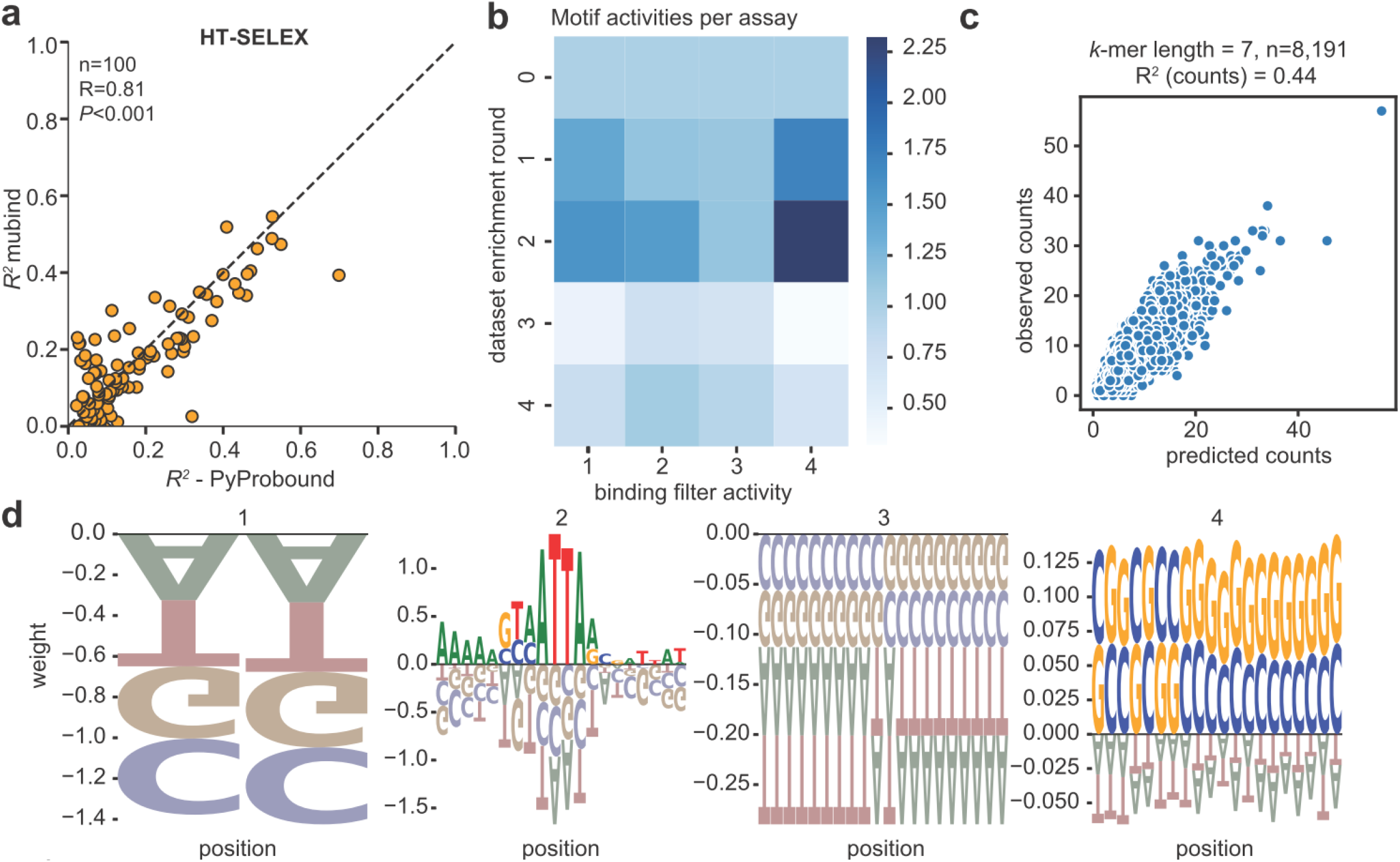
*muBind* benchmark in bulk genomics. (**a**) Agreement between model fits in 100 HT-SELEX datasets using *PyProbound* (*x*-axis) and *muBind* (*y*-axis). Dots display the final *R*^*2*^ for data fits between read count observations and predictions. Details on model selection are provided in the **Methods** section. (**b**) Activities Layer tensor for dataset `HOXB3_ES0_TACATT20NTAC_0`. Colors indicate the relative contribution of learned motifs in each enrichment round. (**c**) visualization of observed versus predicted *k*-mer counts in HOXB3 HT-SELEX. *K*-mers are pooled across all DNA reads listed in all enrichment rounds. (**d**) Binding Layer learned motifs (related to *b*). The learned motif number 2 is associated with a homeodomain transcription factor binding mode, whereas motifs 1, 3, and 4 are associated with likely sequencing biases in the dataset. The motif number 1 is a dinucleotide bias filter.

**Supplementary Figure 2.**
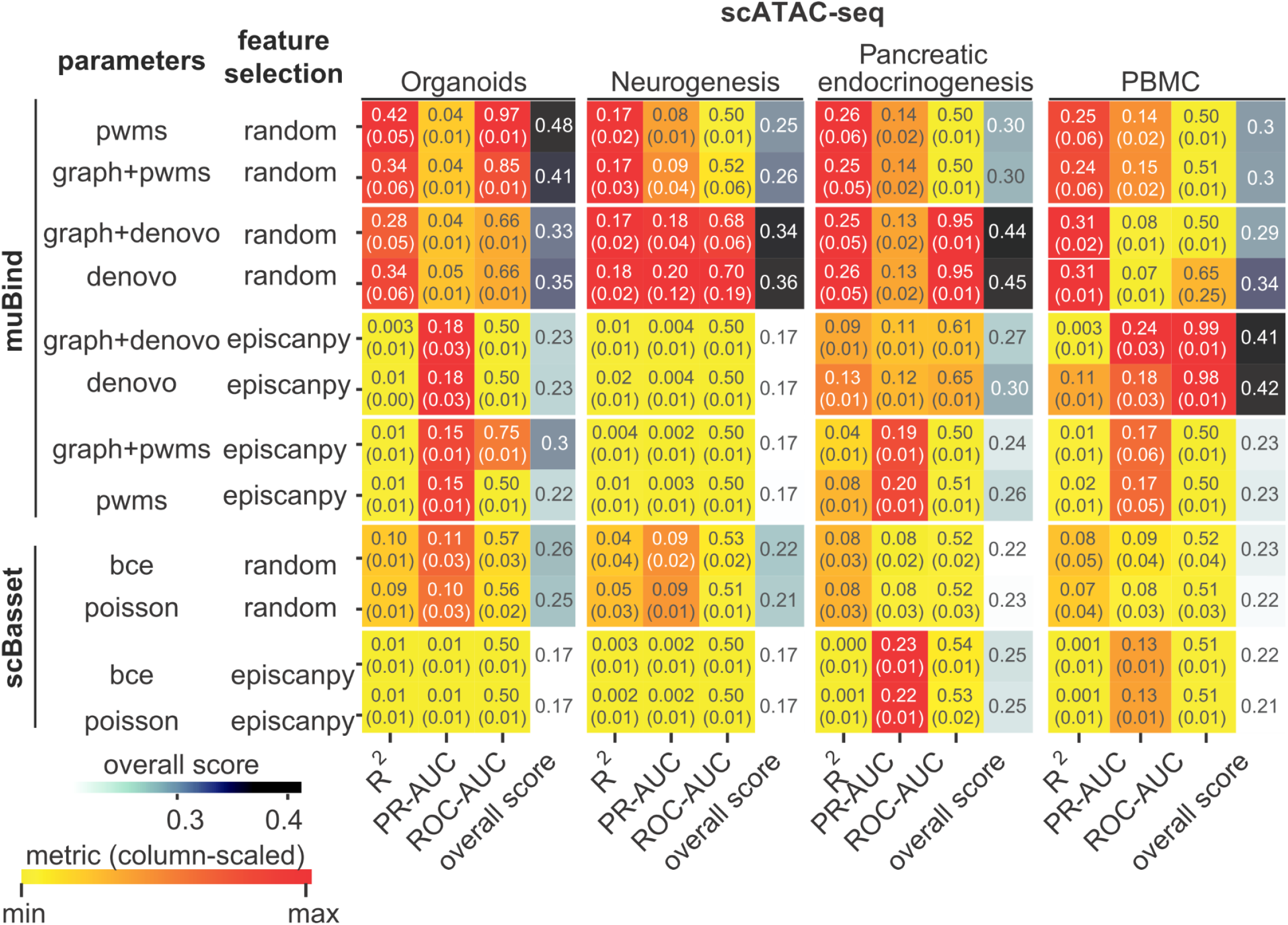
*muBind* accurately predicts genomics counts in single-cell genomics. Comparison of *muBind* models in testing data (n=1,000 cells and 15,000 features, five-fold), using classification (ROC-AUC, PR-AUC, multi-label) and quantitative metrics (*R*^*2*^). Combinations of models with a Graph Layer (*graph*), *de novo* learning of binding filters (*de novo*), and preloaded binding filters that are refined (*pwms*) are shown. The overall score is the mean of each row values, 0-1 normalized per column. Feature selection of features in each dataset is done via random selection (*random*) or with epiScanpy overall accessibility function (*episcanpy*). *scBasset* is tested using Binary Cross Entropy (bce), or Poisson (*poisson*). Value in parenthesis indicates standard deviation of testing metrics (five-fold). Further in Methods.

**Supplementary Figure 3.**
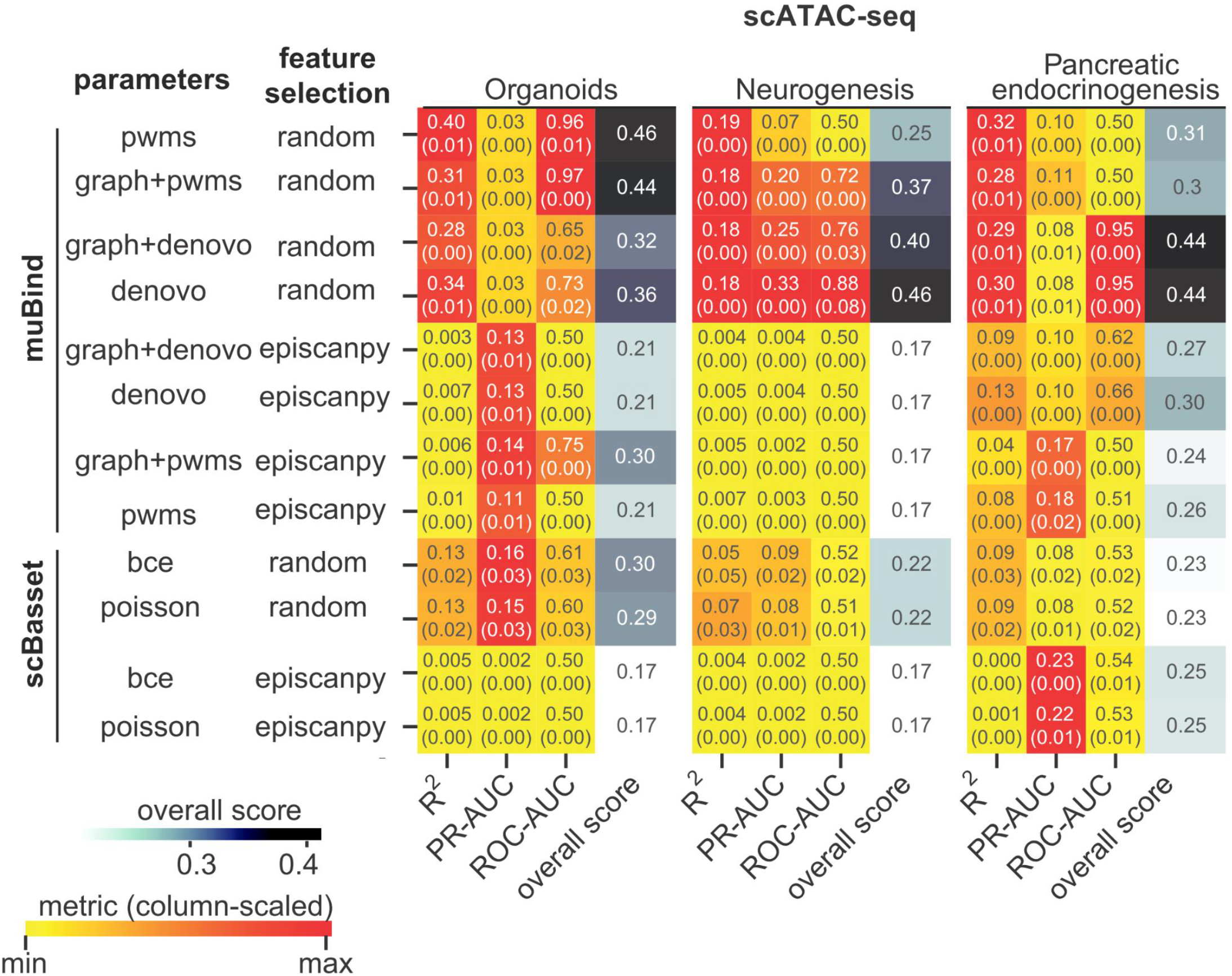
Training metrics for Organoids, Neurogenesis, and Pancreatic endocrinogenesis dataset. Captions as in **Supp Fig 2**

**Supplementary Figure 4.**
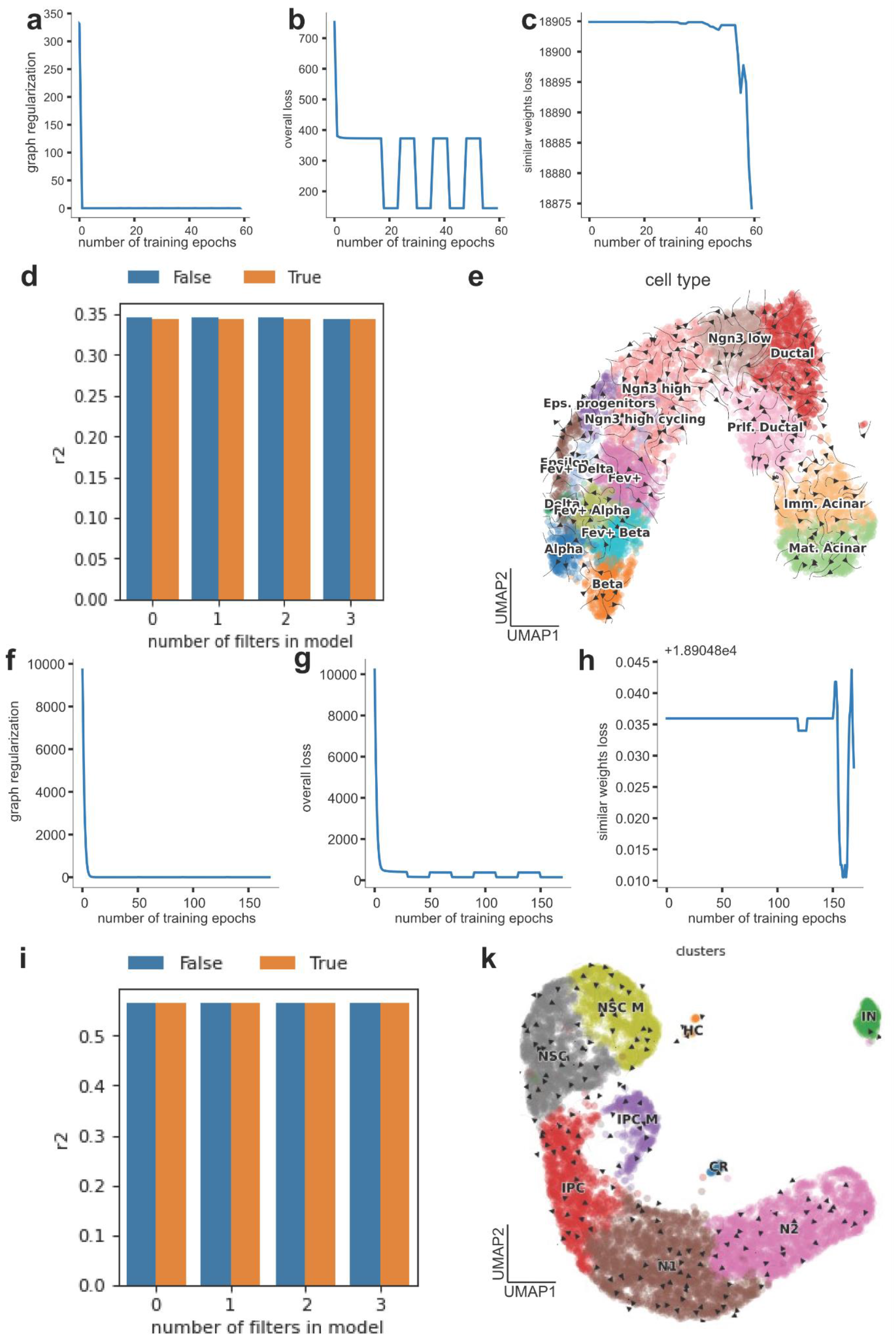
*muBind* losses and scVelo velocity stream without RNA-priors. (*related to Fig 4*) (**a**) Graph regularization over epochs in pancreatic endocrinogenesis. (**b**) Sum of all losses over epochs. Step-wise patterns are associated with weight freezing and unfreezing of filters in the binding layer. (**c**) filter similarities loss over epochs. (**d**) Quantitative performance of PWMs-based *muBind* model, with (True) and without (False) a Graph layer, versus the number of filter groups sequentially optimized. Each value indicates the final prediction R2s values after 50 epochs of training (**e**) UMAP of pancreatiic endocrinogenesis, using Graph Layer learned *D* values to construct a velocity graph, without additional RNA velocity priors from scVelo. (*related to Fig 5*) (**f** to **g**) Equivalent to **a-e** in the mouse neurogenesis dataset.

**Supplementary Figure 5.**
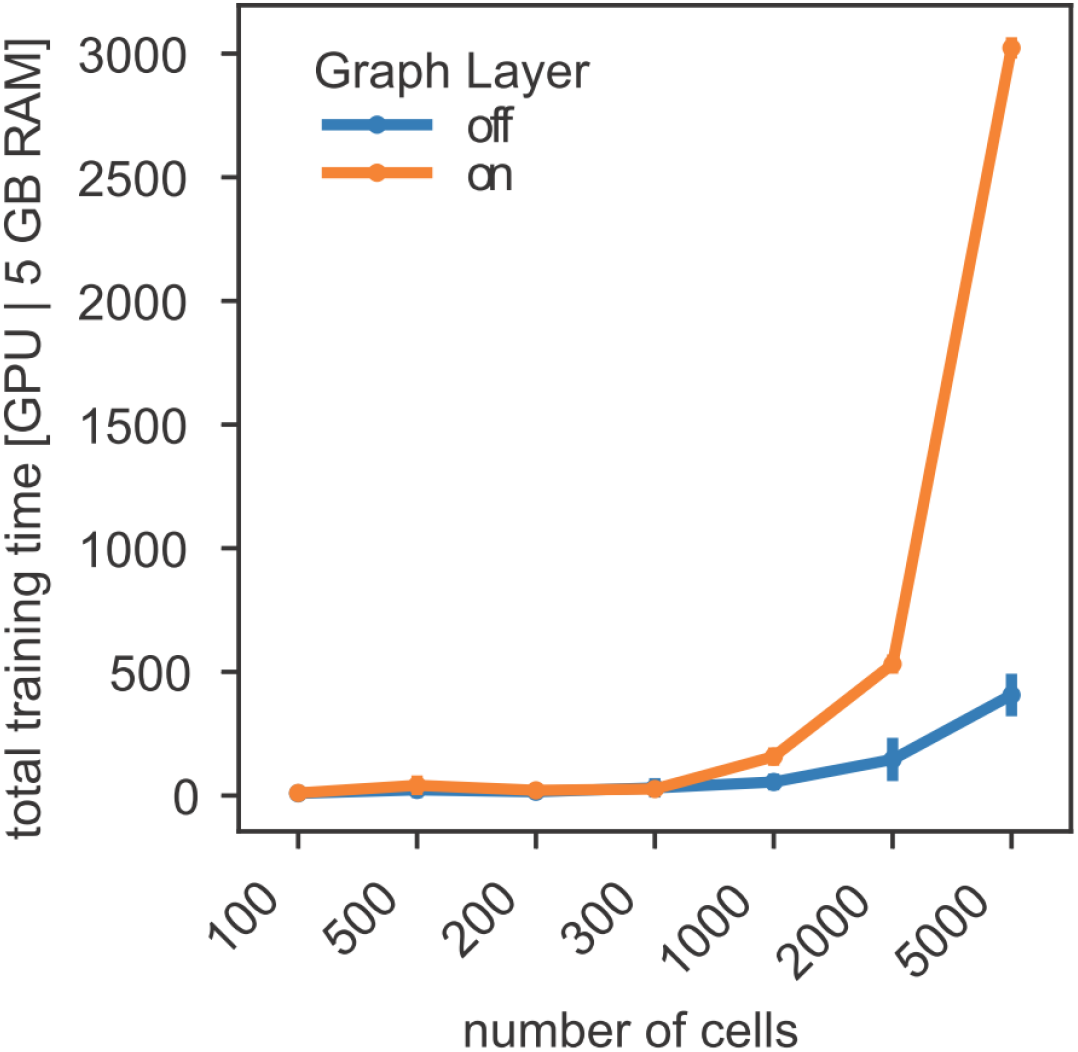
*muBind* running time in single-cell benchmark with and without Graph Layer. Number of features per dataset is set three times the number of cells e.g. 5,000 cells = 15,000 features.

**Supplementary Figure 6.**
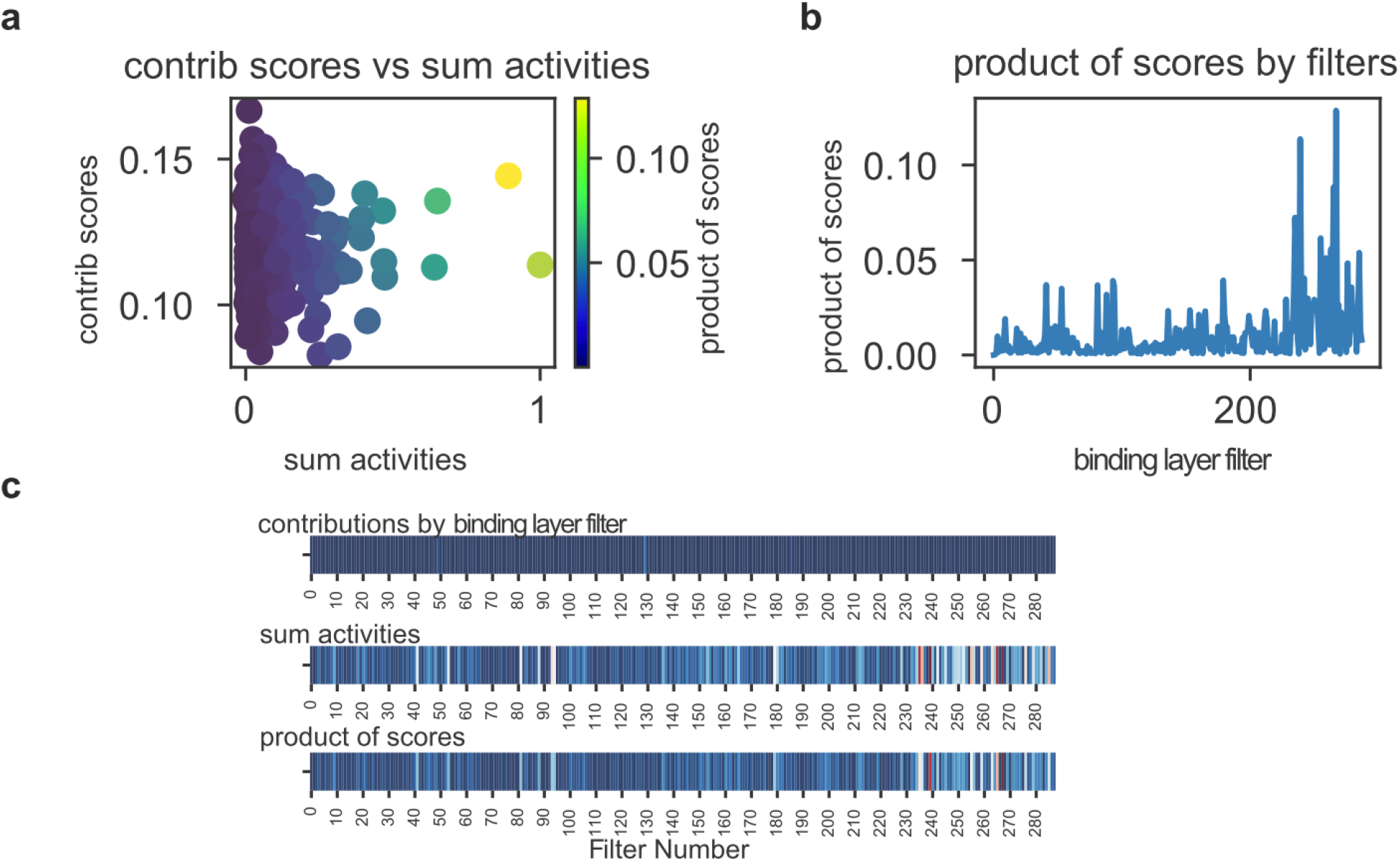
graphs contributions in pancreatic endocrinogenesis scATAC-seq. **(a)** scatter plot visualizing graph contribution scores per filter (y-axis) versus the sum of the absolute value of filter activities (x-axis). Color highlights the product of x and y values. **(b)** product of panel *a* values across the binding layer (n=288). Filters with strong interactions with the graph weights are distributed in different sections of it (peaks). **(c)** similar to *b* showing x and y axes of panel a, and three stacked heatmaps. Values of the top two rows are multiplied to generate the last row value. For details, see **Methods**.

## References

1. Spitz, F. & Furlong, E. E. M. Transcription factors: from enhancer binding to developmental control. Nat. Rev. Genet. 13, 613–626 (2012).

2. Buenrostro, J. D., Wu, B., Chang, H. Y. & Greenleaf, W. J. ATAC-seq: A Method for Assaying Chromatin Accessibility Genome-Wide. Curr. Protoc. Mol. Biol. 109, 21.29.1–21.29.9 (2015).

3. Grosselin, K. et al. High-throughput single-cell ChIP-seq identifies heterogeneity of chromatin states in breast cancer. Nat. Genet. 51, 1060–1066 (2019).

4. Liu, H. et al. DNA methylation atlas of the mouse brain at single-cell resolution. Nature 598, 120–128 (2021).

5. Novakovsky, G., Dexter, N., Libbrecht, M. W., Wasserman, W. W. & Mostafavi, S. Obtaining genetics insights from deep learning via explainable artificial intelligence. Nat. Rev. Genet. 24, 125–137 (2023).

6. Stormo, G. D. & Zhao, Y. Determining the specificity of protein-DNA interactions. Nat. Rev. Genet. 11, 751–760 (2010).

7. Rohs, R. et al. Origins of specificity in protein-DNA recognition. Annu. Rev. Biochem. 79, 233–269 (2010).

8. Rastogi, C. et al. Accurate and sensitive quantification of protein-DNA binding affinity. Proc. Natl. Acad. Sci. U. S. A. 115, E3692–E3701 (2018).

9. Jain, D., Baldi, S., Zabel, A., Straub, T. & Becker, P. B. Active promoters give rise to false positive ‘Phantom Peaks’ in ChIP-seq experiments. Nucleic Acids Res. 43, 6959–6968 (2015).

10. Yuan, H. & Kelley, D. R. scBasset: sequence-based modeling of single-cell ATAC-seq using convolutional neural networks. Nat. Methods 19, 1088–1096 (2022).

11. Avsec, Ž. et al. Base-resolution models of transcription-factor binding reveal soft motif syntax. Nat. Genet. 53, 354– 366 (2021).

12. Rube, H. T. et al. Prediction of protein-ligand binding affinity from sequencing data with interpretable machine learning. Nat. Biotechnol. (2022) doi:10.1038/s41587-022-01307-0.

13. Chen, H. et al. Assessment of computational methods for the analysis of single-cell ATAC-seq data. Genome Biol. 20, 241 (2019).

14. Haghverdi, L., Büttner, M., Wolf, F. A., Buettner, F. & Theis, F. J. Diffusion pseudotime robustly reconstructs lineage branching. Nat. Methods 13, 845–848 (2016).

15. Weiler, P., Lange, M., Klein, M., Pe’er, D. & Theis, F. CellRank 2: unified fate mapping in multiview single-cell data. Nat. Methods (2024) doi:10.1038/s41592-024-02303-9.

16. Li, H. et al. Inferring transcription factor regulatory networks from single-cell ATAC-seq data based on graph neural networks. Nature Machine Intelligence 4, 389–400 (2022).

17. Fleck, J. S. et al. Inferring and perturbing cell fate regulomes in human brain organoids. Nature 621, 365–372 (2023).

18. Kamimoto, K. et al. Dissecting cell identity via network inference and in silico gene perturbation. Nature 614, 742– 751 (2023).

19. Bravo González-Blas, C. et al. SCENIC+: single-cell multiomic inference of enhancers and gene regulatory networks. Nat. Methods 20, 1355–1367 (2023).

20. Wolf, F. A., Angerer, P. & Theis, F. J. SCANPY: Large-scale single-cell gene expression data analysis. Genome Biol. 19, 15 (2018).

21. Moon, K. R. et al. Manifold learning-based methods for analyzing single-cell RNA-sequencing data. Current Opinion in Systems Biology 7, 36–46 (2018).

22. Vierstra, J. et al. Global reference mapping of human transcription factor footprints. Nature 583, 729–736 (2020).

23. Virshup, I. et al. The scverse project provides a computational ecosystem for single-cell omics data analysis. Nat. Biotechnol. 41, 604–606 (2023).

24. Hochgerner, H., Zeisel, A., Lönnerberg, P. & Linnarsson, S. Conserved properties of dentate gyrus neurogenesis across postnatal development revealed by single-cell RNA sequencing. Nat. Neurosci. 21, 290–299 (2018).

25. Satpathy, A. T. et al. Massively parallel single-cell chromatin landscapes of human immune cell development and intratumoral T cell exhaustion. Nat. Biotechnol. 37, 925–936 (2019).

26. Noack, F. et al. Multimodal profiling of the transcriptional regulatory landscape of the developing mouse cortex identifies Neurog2 as a key epigenome remodeler. Nat. Neurosci. 25, 154–167 (2022).

27. Klein, D. et al. Mapping cells through time and space with moscot. bioRxiv 2023.05.11.540374 (2023) doi:10.1101/2023.05.11.540374.

28. Fischer, D. S. et al. Sfaira accelerates data and model reuse in single cell genomics. Genome Biol. 22, 248 (2021).

29. Danese, A. et al. EpiScanpy: integrated single-cell epigenomic analysis. Nat. Commun. 12, 5228 (2021).

30. Martens, L. D., Fischer, D. S., Yépez, V. A., Theis, F. J. & Gagneur, J. Modeling fragment counts improves single-cell ATAC-seq analysis. Nat. Methods 21, 28–31 (2024).

31. Tedesco, M. et al. Chromatin Velocity reveals epigenetic dynamics by single-cell profiling of heterochromatin and euchromatin. Nat. Biotechnol. 40, 235–244 (2022).

32. Li, C., Virgilio, M. C., Collins, K. L. & Welch, J. D. Multi-omic single-cell velocity models epigenome-transcriptome interactions and improves cell fate prediction. Nat. Biotechnol. 41, 387–398 (2023).

33. Bergen, V., Lange, M., Peidli, S., Wolf, F. A. & Theis, F. J. Generalizing RNA velocity to transient cell states through dynamical modeling. Nat. Biotechnol. 1–7 (2020).

34. Seymour, P. A. Sox9: a master regulator of the pancreatic program. Rev. Diabet. Stud. 11, 51–83 (2014).

35. Piper, K. et al. Novel SOX9 expression during human pancreas development correlates to abnormalities in Campomelic dysplasia. Mech. Dev. 116, 223–226 (2002).

36. Hale, M. A. et al. The nuclear hormone receptor family member NR5A2 controls aspects of multipotent progenitor cell formation and acinar differentiation during pancreatic organogenesis. Development 141, 3123–3133 (2014).

37. ENCODE Project Consortium et al. Expanded encyclopaedias of DNA elements in the human and mouse genomes. sNature 583, 699–710 (2020).

38. Han, H. et al. TRRUST v2: an expanded reference database of human and mouse transcriptional regulatory interactions. Nucleic Acids Res. 46, D380–D386 (2018).

39. Hasenpusch-Theil, K. et al. Gli3 controls the onset of cortical neurogenesis by regulating the radial glial cell cycle through Cdk6 expression. Development 145, (2018).

40. Wang, J. et al. FOXG1 Contributes Adult Hippocampal Neurogenesis in Mice. Int. J. Mol. Sci. 23, (2022).

41. Salih, D. A. M. et al. FoxO6 regulates memory consolidation and synaptic function. Genes Dev. 26, 2780–2801 (2012).

42. Hou, P.-S., hAilín, D.Ó., Vogel, T. & Hanashima, C. Transcription and Beyond: Delineating FOXG1 Function in Cortical Development and Disorders. Front. Cell. Neurosci. 14, 35 (2020).

43. Prajapati, R. S., Hintze, M. & Streit, A. PRDM1 controls the sequential activation of neural, neural crest and sensory progenitor determinants. Development 146, (2019).

44. Baizabal, J.-M. et al. The Epigenetic State of PRDM16-Regulated Enhancers in Radial Glia Controls Cortical Neuron Position. Neuron 98, 945–962.e8 (2018).

45. He, L. et al. PRDM16 regulates a temporal transcriptional program to promote progression of cortical neural progenitors. Development 148, (2021).

46. Burdziak, C. et al. scKINETICS: inference of regulatory velocity with single-cell transcriptomics data. Bioinformatics 39, i394–i403 (2023).

47. Aliee, H. et al. Sparsity in Continuous-Depth Neural Networks. arXiv [cs.LG] (2022).

48. Lopez, R., Regier, J., Cole, M. B., Jordan, M. I. & Yosef, N. Deep generative modeling for single-cell transcriptomics. Nat. Methods 15, 1053–1058 (2018).

49. Luecken, M. D. et al. Benchmarking atlas-level data integration in single-cell genomics. Nat. Methods 19, 41–50 (2022).

50. Haghverdi, L., Lun, A. T. L., Morgan, M. D. & Marioni, J. C. Batch effects in single-cell RNA-sequencing data are corrected by matching mutual nearest neighbors. Nat. Biotechnol. 36, 421–427 (2018).

51. Jolma, A. et al. DNA-binding specificities of human transcription factors. Cell 152, 327–339 (2013).

